# Machine Classification of Methylomes in Cancer

**DOI:** 10.1101/2020.04.04.025155

**Authors:** Isabelle Newsham, Marcin Sendera, SriGanesh Jammula, Rebecca Fitzgerald, Charles Massie, Shamith A. Samarajiwa

**Affiliations:** MRC Cancer Unit, Cambridge Biomedical Campus, Box 197, University of Cambridge, Cambridge, CB2 0XZ, UK; Department of Oncology, Cambridge Biomedical Campus, Box 197, University of Cambridge, CB2 0XZ, UK; CRUK Cambridge Institute, University of Cambridge, Robinson Way, CB2 ORE, UK

## Abstract

Cancer remains a leading cause of morbidity and mortality worldwide. Its evolutionary nature and resultant complex interactions with the tumour micro-environment and the host immune system engender heterogeneity, make developing interventions difficult. Usually detected at the advanced stages of disease, metastatic cancer accounts for 90% of cancer-associated deaths. Therefore early detection of cancer, combined with current therapies, would have a significant impact on survival and treatment of this insidious disease. Epigenetic changes such as DNA methylation are some of the early events in carcinogenesis. Here, we report on a machine learning model that can classify 13 types of cancer as well as non-cancer tissue samples using only DNA methylome data, with an accuracy of 98.2%. We utilise the features identified by this model to develop a robust deep neural network that can generalise to independent data sets. We also demonstrate that the methylation associated genomic loci detected by the classifier are associated with genes involved in cancer, providing insights into the epigenomic regulation of carcinogenesis.

## 1 Introduction

Each of our cells contain a single identical genome, incorporating the information necessary to specify and maintain our characteristics. In contrast, each cell will exhibit multiple epigenomes that change during different cellular states and over the passage of time. These epigenomes consist of a collection of reversible chromatin interactions and modifications that are heritable and do not change the DNA sequence. Histone variations, post translational modifications of the amino terminal tails of histone proteins, nucleosomal positioning, non-coding RNA interactions and covalent modification of DNA are some of the factors that contribute to epigenomic change. Notably, covalent methylation of DNA is one such reversible chemical modification with many functional consequences, and evidence for its role in genome maintenance and the regulation of gene expression has accumulated in the last few decades [1].

Most commonly, methylation occurs at CpG residues, modifying Cytosine nucleotides to 5-methylcytosine (5mC). In humans, large stretches of CpGs (regions greater than 500 bp with more than 50% CpG content), known as CpG islands (CGI), contribute to gene regulatory promoter elements of over 72% of protein coding genes [2]. These CGI regions are usually protected from methylation, however their aberrant methylation is observed in many cancers. Consequently, epigenetic modifications are some of the earliest neoplastic events associated with carcinogenesis [3, 4].

Regulatory proteins with affinity to methylated regions cooperate with chromatin modifiers to establish focally silenced states. Thus, DNA methylation inhibits transcription factor (TF) recognition, affinity and accessibility of its DNA motifs [5]. CpG island promoter hypermethylation of tumour suppressor genes is an early neoplastic event in many tumours [6, 7, 8, 9]. Epigenetic instability can also be induced by a mechanism known as the CpG island methylator phenotype (CIMP). Initially identified in colorectal cancer, CIMP is characterised by promoter CpG island hypermethylation of the mismatch repair gene, MLH1, resulting in its transcriptional inactivation. This loss of MLH1 activity leads to microsatellite instability (MSI), which causes length alterations within simple microsatellite repeats leading to an increase in the rate of carcinogenic progression [8, 10]. Conversely, global DNA hypomethylation can lead to chromosomal instability, activation of oncogenes and latent retro-transposons that promote carcinogenesis [11]. Hypomethylation is seen in many cancer types, including cervical, prostate, hepatocellular, breast, brain and leukaemia [12, 13, 14, 15].

These hyper- and hypo-methylation patterns can serve as cancer associated signals and prognostic biomarkers. Computational methods that detect these complex neoplastic methylation patterns can assist in cancer early detection, diagnosis and screening. We developed both binary and multiclass machine learning models to identify multiple cancer types from non-cancerous tissue samples. Our method is robust, generalisable and interpretable and demonstrates high predictive accuracy. Due to its ease of use and adaptability, it could be used in screening of multiple cancers. It is also extensible and can be applied in potential applications of early detection of cancer.

## 2 Results

**Overview** We utilised machine learning approaches to identify cancer specific changes from normal tissue specific methylation. DNA methylation microarray data from 13 cancer types and corresponding normal tissues were utilised. Data was extracted, cleaned and processed as described in the Methods. Illumina Infinium array-based methylome data was used in this study. This microarray platform detects methylation at given CpG locations, using a pair of methylated and unmethylated probes. In this study, we created and evaluated four different model types: logistic regression, support vector machines (SVM), gradient boosted decision trees (XGBoost), and a deep neural network (DNN). See Figure 1 for a visual overview. For the first three model types, both binary and multiclass classification models were created.

**Figure 1:**
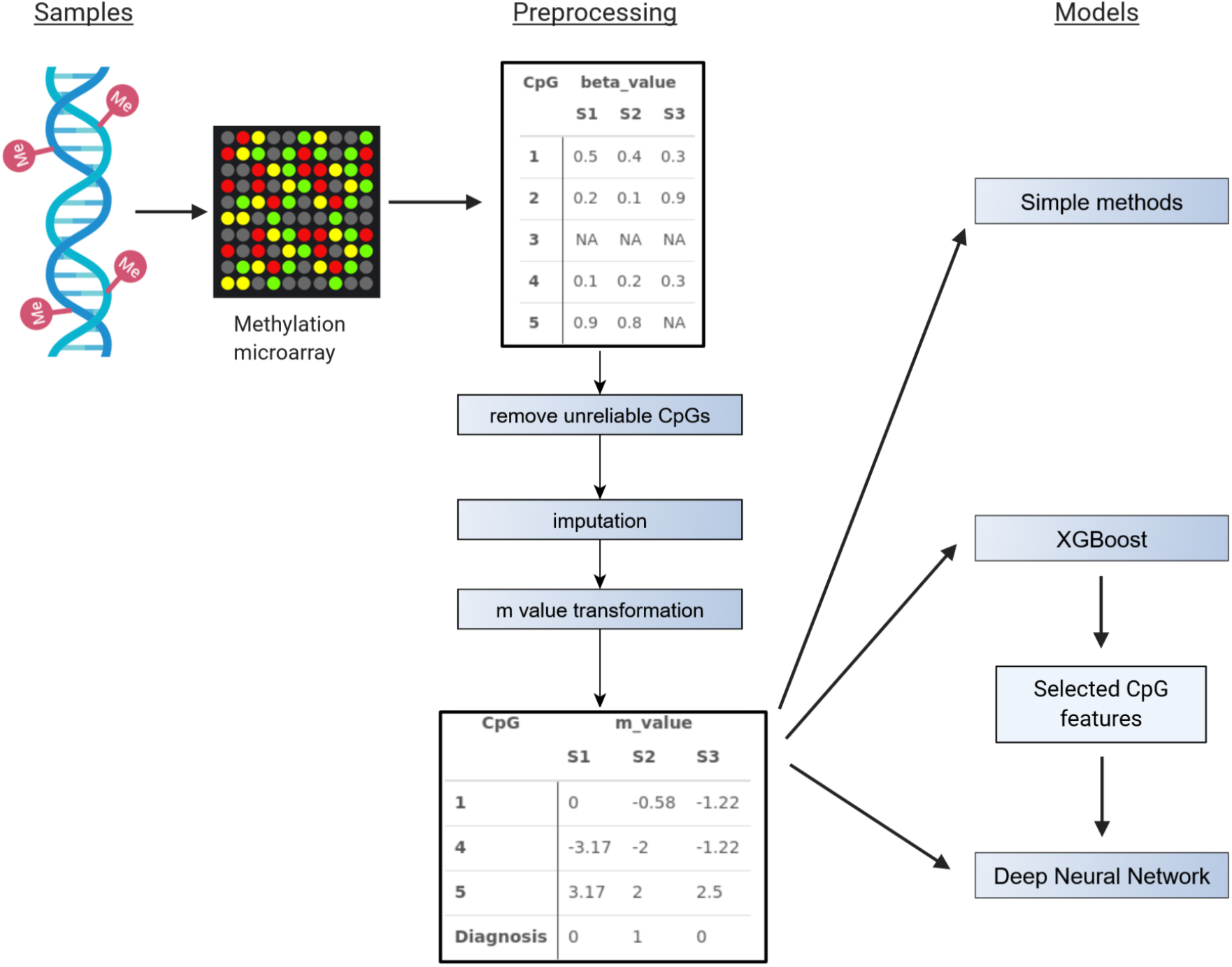
Overview of method. DNA methylation microarray data from 13 cancer types and corresponding normal tissues were collected from TCGA. Data were preprocessed by removing unreliable features (CpGs), imputating missing values, and transforming to M-values. Then three types of models were trained: Simple methods (logistic regression and support vector machines), XGBoost, and a Deep Neural Network.

### 2.1 Training, testing and model evaluation

Here, we present the results from the XGBoost and deep neural network models. The logistic regression and SVM models performed surprisingly well, but on the whole were less robust, prone to over-fitting and did not outperform the XGBoost or the deep neural network (see Supplementary Tables S1 and S2).

**Binary classification of individual tumour and normal tissues** Gradient boosted decision tree models are an iterative ensemble machine learning approach. Boosting trains a succession of models with each new model being trained to correct the errors made by the previous ones. During this process, models are accrued until no further improvements can be made. XGBoost is a python library with an underlying C++ code-base that implements gradient boosted decision trees [16].

We trained 13 binary XGBoost models, one for each cancer type. Each model learns to classify between cancer and normal samples for its tissue type. Overall there was good performance on the test set, with 5 out of 13 models achieving a perfect test set performance (colon adenocarcinoma, kidney renal clear cell carcinoma, lung adenocarcinoma, lung squamous cell carcinoma, and uterine corpus endometrial carcinoma). Across all models, the average accuracy was 0.987 and the average Matthews Correlation Coefficient (MCC, a performance measure unaffected by severe class imbalance) was 0.919, demonstrating that the models can accurately classify cancer and normal samples. Figure 2 shows the confusion matrices for the best and worst performing models, and the Area Under the Curve (AUC) of ROC curves and precision-recall curves for all models. Percentage performance metrics for all binary models can be found in Supplementary Tables S3. A key issue with these binary models is the major class imbalance. The average fraction of normal samples in a tissue type is 0.135, which reveals why the average MCC is considerably lower than the average accuracy. In addition, the lowest performing model, esophageal carcinoma (ESCA), with an accuracy of 0.961 and MCC of 0.693, is the tissue type with the lowest number of normal samples, of which there were only 16. This lack of data possibly contributed towards its worse performance.

**Figure 2:**
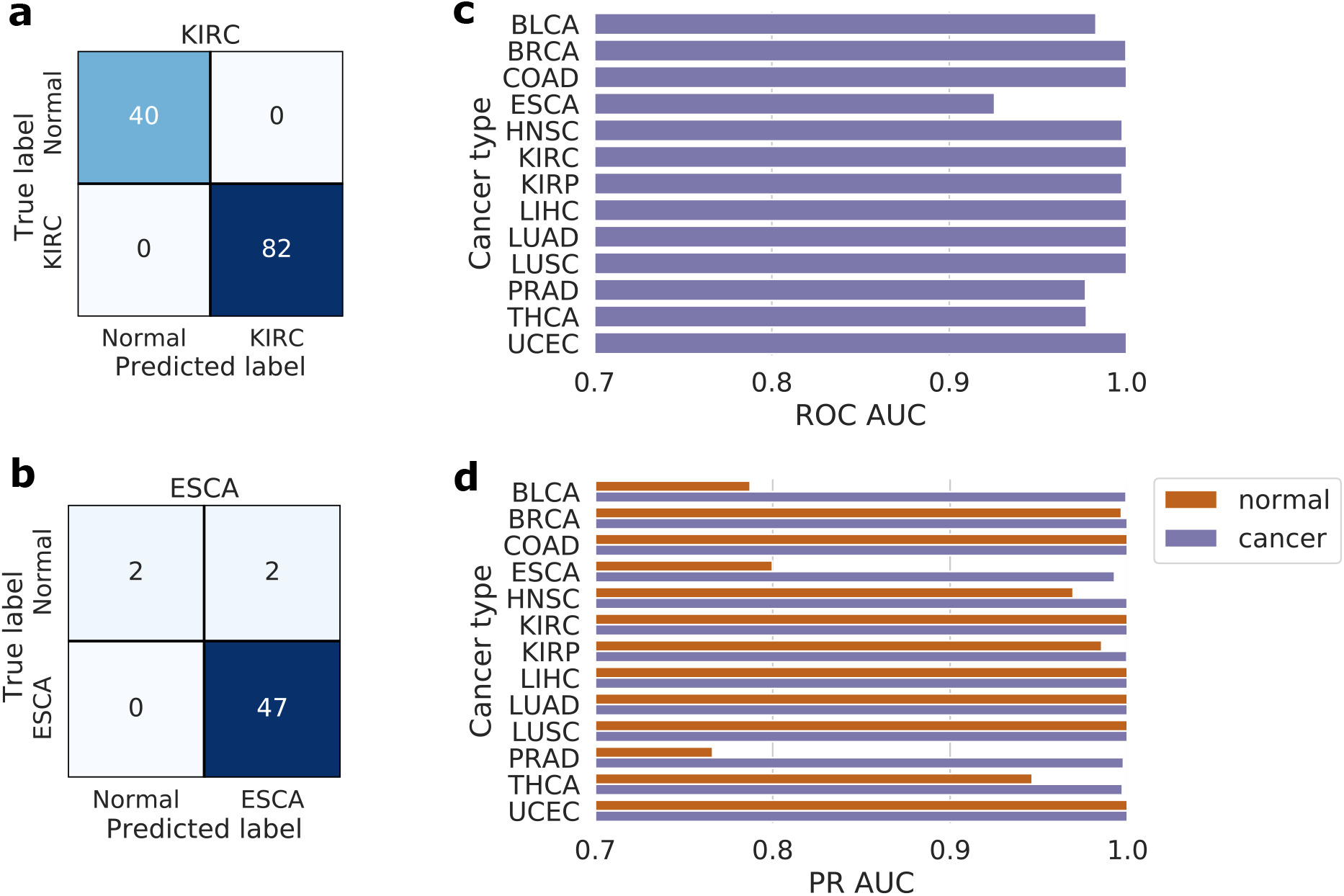
Performance of the binary XGBoost models on the TCGA test set. **a** and **b** Confusion matrices of the best (KIRC) and worst (ESCA) performing binary XGBoost models. **c** AUC of the ROC curves for all binary XGBoost models. **d** AUC of the Precision Recall (PR) curves for both cancer and normal classes of all binary XGBoost models. Note that the scales of **c** and **d** start from 0.7.

**Multiclass classification of 13 cancer and normal tissues** Here, we trained a single multiclass XGBoost model on the whole of the training data. There were classes for each of the 13 cancer types and a single normal class, which contained normal samples from every cancer type. The model was now required to learn the differences between 13 tissue types in addition to the differences between cancer and normal tissue samples, making it a more difficult task than the previous binary classification. However, there was no longer a large class imbalance due to pooling of the normal samples together. As shown in Figure 3, the performance of the test set was very good for all classes. The model can discriminate each of the 13 cancer types and normal samples with a high degree of accuracy. The overall accuracy was 0.982 and the overall MCC was 0.980, see Table 1 for the detailed metrics.

**Figure 3:**
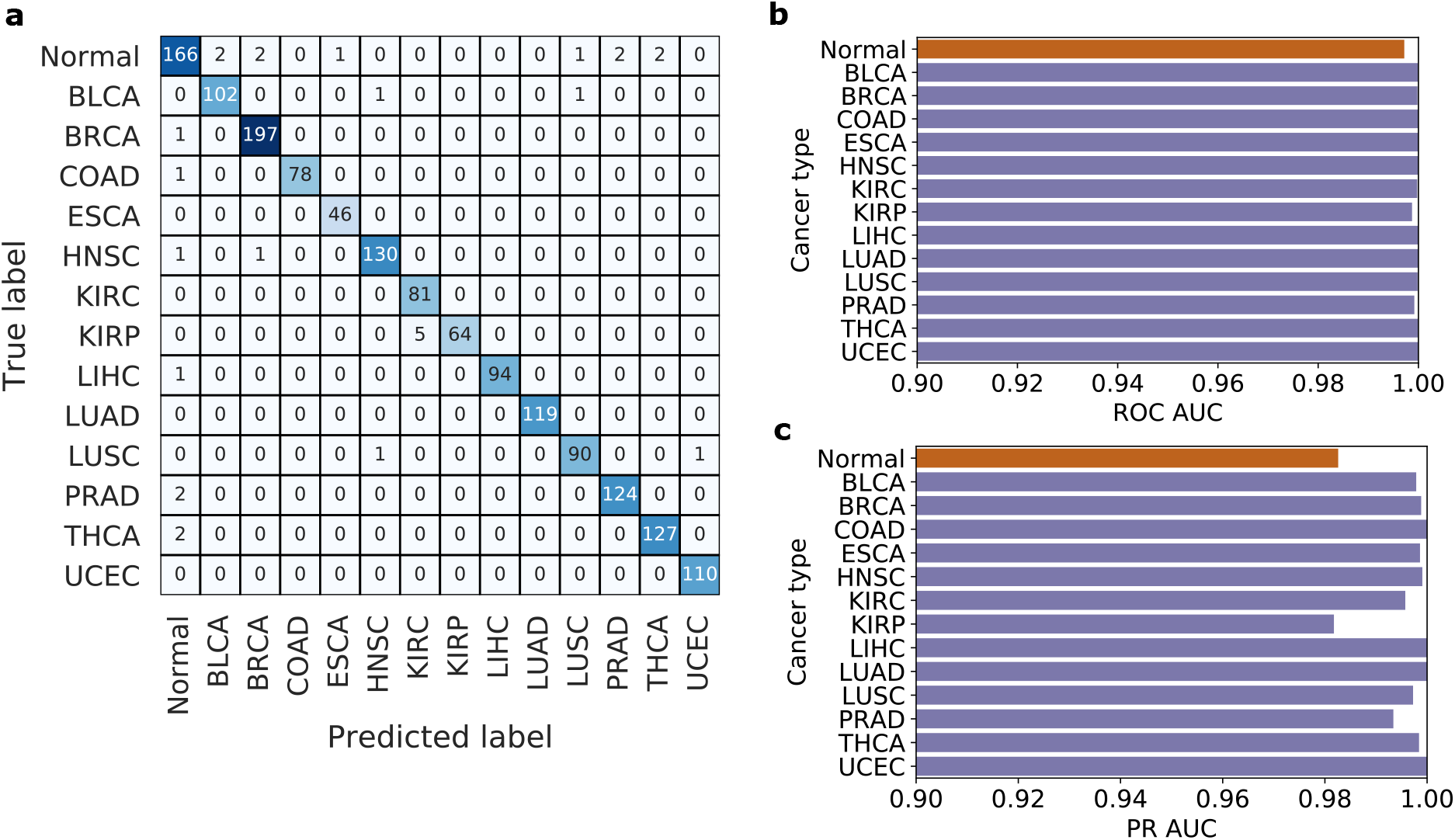
Performance of the multiclass XGBoost model on the TCGA test set. **a** shows the confusion matrix, **b** shows the AUC of the ROC curves for each class, and **c** shows the AUC of the Precision Recall (PR) curves for each class. The colour orange denotes normal and purple denotes cancer. Note that the scales of **b** and **c** start from 0.9.

**Table 1:** Performance metrics for the multiclass XGBoost model

|  | Normal | BLCA | BRCA | COAD | ESCA | HNSC | KIRC | KIRP | LIHC | LUAD | LUSC | PRAD | THCA | UCEC |
| --- | --- | --- | --- | --- | --- | --- | --- | --- | --- | --- | --- | --- | --- | --- |
| Precision | 0.954 | 0.981 | 0.985 | 1.0 | 0.979 | 0.985 | 0.942 | 1.0 | 1.0 | 1.0 | 0.978 | 0.984 | 0.984 | 0.991 |
| Recall | 0.943 | 0.981 | 0.995 | 0.987 | 1.0 | 0.985 | 1.0 | 0.928 | 0.989 | 1.0 | 0.978 | 0.984 | 0.984 | 1.0 |
| $F_1$ | 0.949 | 0.981 | 0.99 | 0.994 | 0.989 | 0.985 | 0.97 | 0.962 | 0.995 | 1.0 | 0.978 | 0.984 | 0.984 | 0.995 |
| Accuracy | 0.982 |  |  |  |  |  |  |  |  |  |  |  |  |  |
| MCC | 0.980 |  |  |  |  |  |  |  |  |  |  |  |  |  |

### 2.2 Performance on independent data

To determine the robustness of our models, we tested our XGBoost models on a number of independent data sets, representing different cancer types. These were more heterogeneous than the TCGA data used for training. Two data sets included some adenoma samples (COAD and THCA), one data set consisted of samples from early-stage tumours, some of which were later shown to recur (LIHC), and one data set included some HPV positive samples (HNSC).

The data sets also came from seven different countries, viz., Iceland (BRCA), USA (COAD), Australia (ESCA), UK (HNSC), China (LIHC and KIRC), Canada (PRAD), and Brazil (THCA).

**Binary models** When these independent data sets were tested, most of the binary XGBoost models (trained on TCGA data) performed well, illustrated by Figure 4. The highest performing model was the breast cancer (BRCA) model, with a perfect ROC AUC of 1.0, and the lowest performing was the colon adenocarcinoma (COAD) model, with a ROC AUC of 0.758.

**Figure 4:**
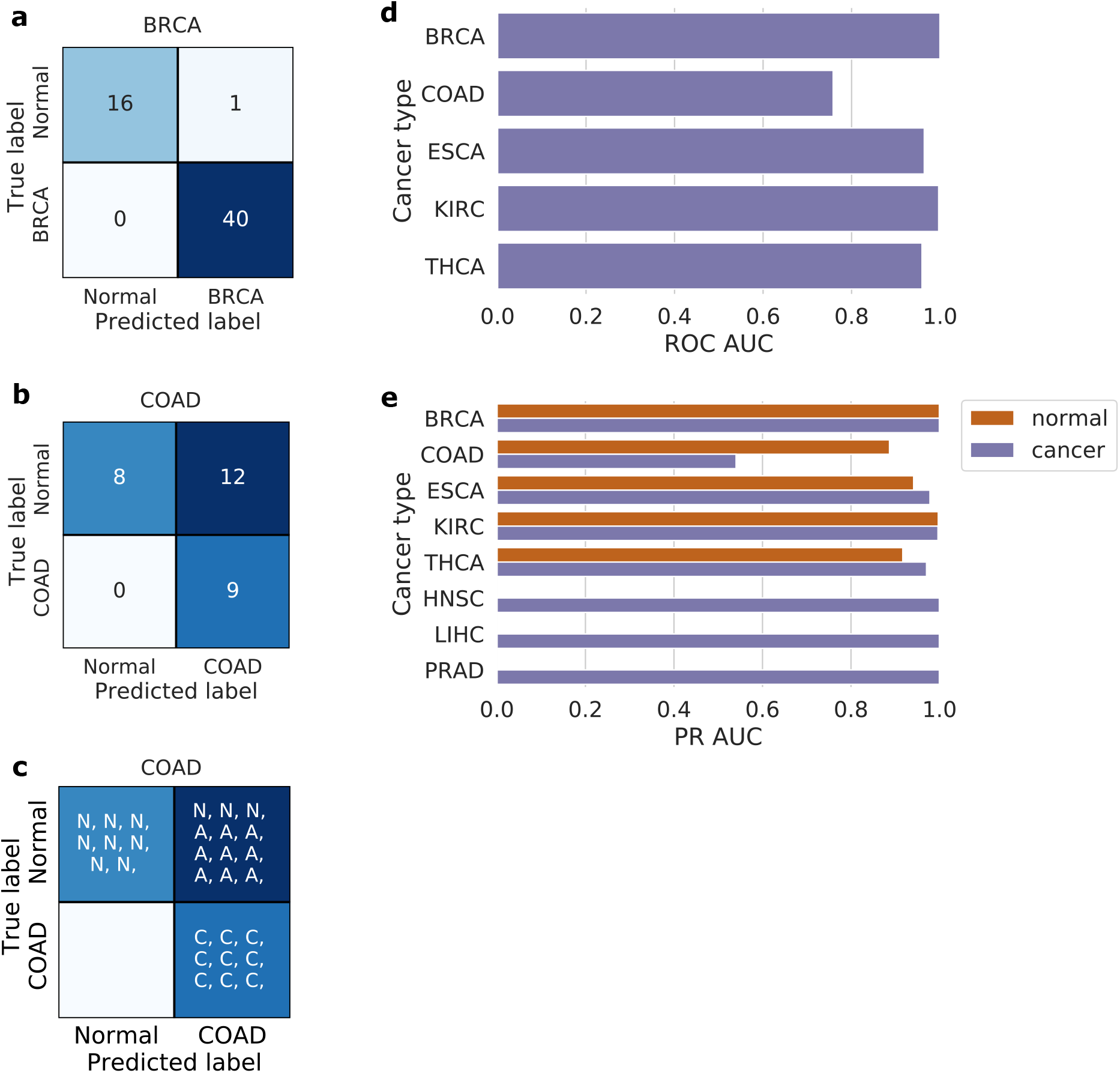
Performance of the binary XGBoost models on independent data sets. **a** and **b** Confusion matrices of the best (BRCA) and worst (COAD) performing binary XGBoost models on the independent data sets. **c** Detailed confusion matrix for COAD showing the predictions of Normal (N), Adenoma (A), and Cancer (C) samples. **d** AUC of the ROC curves for binary XGBoost models where the independent data set included normal samples. **e** AUC of the Precision Recall (PR) curves for both cancer and normal (where available) classes of binary XGBoost models on the independent data sets.

The confusion matrix of the COAD model in Figure 4**b** shows that it predicted 12 normal samples as cancer. Nine out of these 12 samples are in fact adenomas; benign tumours of glandular origin. A confusion matrix that also shows whether the samples are Normal (N), Adenomas (A), or Carcinomas (C) is shown in Figure 4**c**, illustrating that all adenomas are classified as cancer. This was unexpected, as there were no adenomas in the training data set and instead of randomly classifying them, the model found some cancer associated signal in adenoma samples in the independent data set.

A similar trend was identified in the other independent data set with adenomas. In the thyroid carcinoma (THCA) model, 11 out of 17 adenomas were predicted to be cancer. In detail, it can be seen that all occurrences of ‘follicular adenoma’, and ‘follicular adenoma/Hürthle cell’ were classified as cancer (*n*=8), all ‘lymphocytic thyroiditis’ were classified as normal (*n*=3), and ‘nodular goiter’ was split evenly between the two classes (*n*=6).

**Multiclass models** The results for the multiclass XGBoost model on the independent data, which had an accuracy of 0.653 and MCC of 0.645, can be found in Supplementary Table S5. With the aim of creating a more robust model and improving these results, we designed a feed forward neural network based on our XGBoost model, as shown in Figure 5**a**. See Supplementary Table S4 for its results on the TCGA test set. The results on the independent data sets, which had an accuracy of 0.856 and MCC of 0.834, are shown in Figure 5 and Table 2. The only data set that did not reach an F_1_ score of at least 0.8 (excluding COAD, as it contains adenomas, see above) was head and neck squamous carcinoma (HNSC).

**Figure 5:**
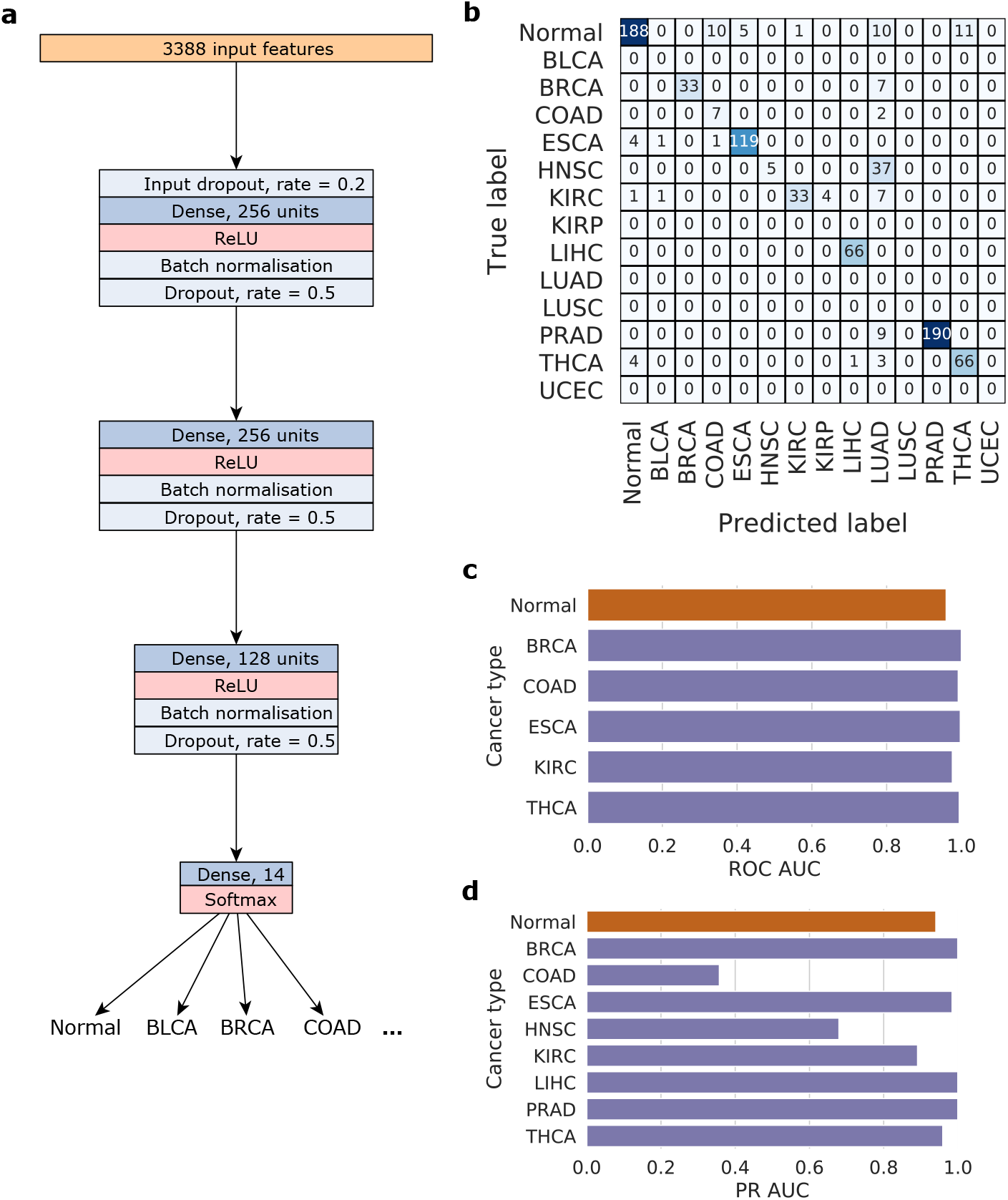
Architecture of the feed forward neural network (**a**) and its performance on all independent data sets. **b** shows the confusion matrix, **c** shows the AUC of the ROC curves for each class where normal samples are available, and **d** shows the AUC of the Precision Recall (PR) curves for each class. The colour orange denotes normal and purple denotes cancer.

**Table 2:** Performance metrics for the multiclass DNN model on independent data

|  | Normal | BRCA | COAD | ESCA | HNSC | KIRC | LIHC | PRAD | THCA |
| --- | --- | --- | --- | --- | --- | --- | --- | --- | --- |
| Precision | 0.954 | 1.0 | 0.389 | 0.96 | 1.0 | 0.971 | 0.985 | 1.0 | 0.857 |
| Recall | 0.836 | 0.825 | 0.778 | 0.952 | 0.119 | 0.717 | 1.0 | 0.955 | 0.892 |
| $F_1$ | 0.891 | 0.904 | 0.519 | 0.956 | 0.213 | 0.825 | 0.992 | 0.977 | 0.874 |
| Accuracy | 0.856 |  |  |  |  |  |  |  |  |
| MCC | 0.834 |  |  |  |  |  |  |  |  |

### 2.3 Biological Feature Interpretability

A key advantage of using an interpretable method such as XGBoost is that the features utilised for classification can be identified. In our case these were the CpG probes with a feature importance of above zero, which we refer to as Probes Contributing to Classification (PCCs). These PCCs can then be mapped to the proximal genes to obtain gene lists. Firstly, we compared our PCCs to probes that were found differentially methylated in a particular cancer type. We chose breast cancer (BRCA), as it has the most number of samples. Surprisingly, only 13 probes are found by both methods, as shown in Figure 6**a**.

**Figure 6:**
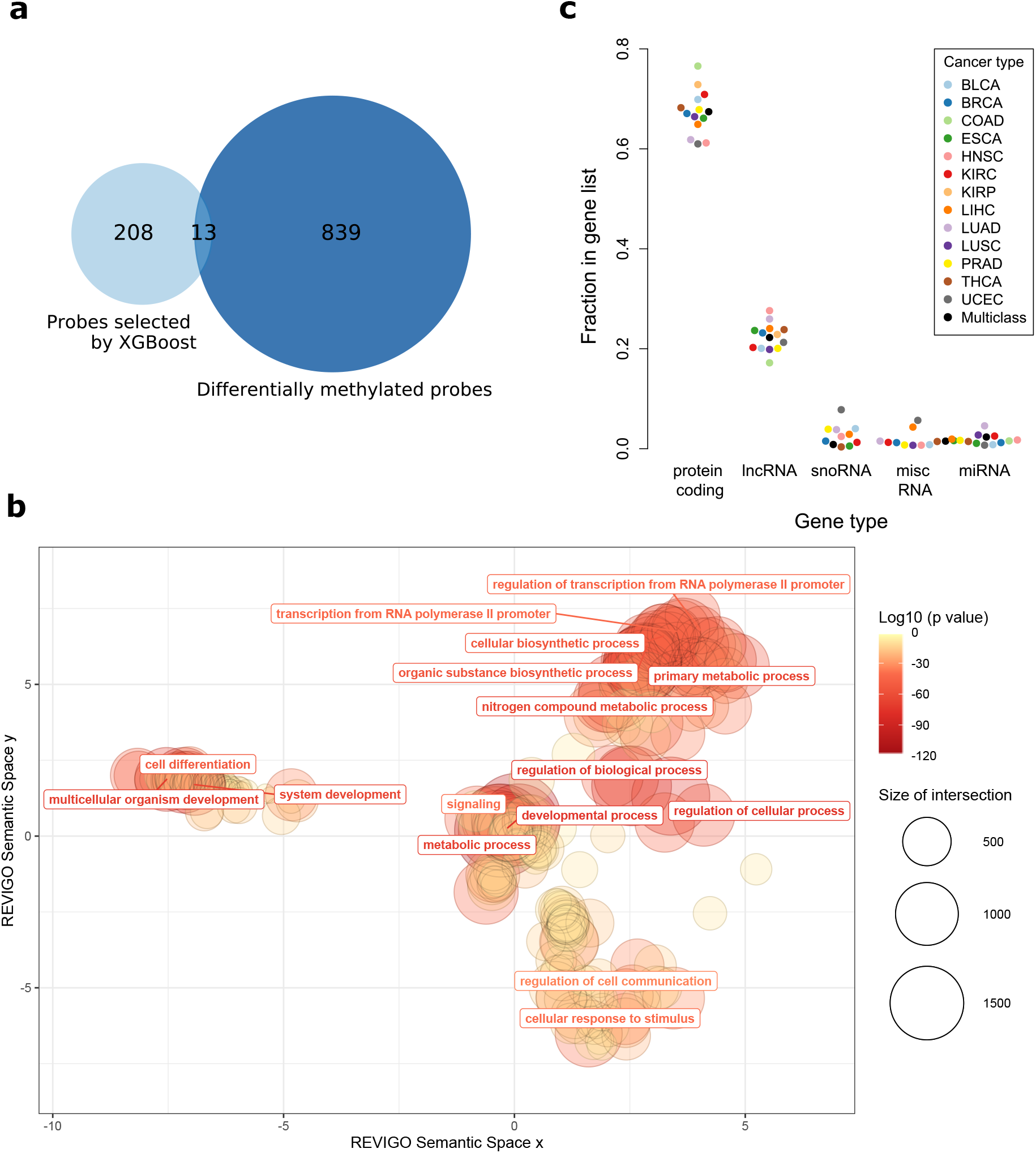
**a** Venn diagram comparing probes selected by the BRCA XGBoost model and probes found through differential methylation analysis, on the TCGA BRCA data. **b** A REVIGO visualisation showing the significant Gene Ontology terms, restricted to the biological process domain. Only a small selection of terms are labelled. **c** The fractions of different gene types in all cancer gene lists, including the multiclass gene list.

Next, we performed functional enrichment analysis on the gene list that was obtained by mapping all multiclass PCCs (the same probes that are the input to the neural network) to proximal genes. A visualisation of the significant Gene Ontology terms, restricted to the biological process ontology, is shown in Figure 6**b**. This shows that our multiclass gene list is enriched in developmental and differentiation, cell communication, metabolic processes, and regulation of transcription.

Finally, we looked into the proportion of protein-coding versus non-coding genes in our gene lists. This is visualised in Figure 6**c** for all individual cancer gene lists and the multiclass gene list. It shows that, as expected, most genes are protein-coding (around 70%). However, the proportion of long non-coding RNA (lncRNA) is surprisingly high for all gene lists (around 25%), which motivated further analysis. As we could not find a comprehensive database of cancer lncRNAs, we used the Pangaea package, which enables text mining of millions of PubMed abstracts [17]. We searched over four million cancer abstracts, and collected abstracts that mentioned at least one of the lncRNAs found in our multiclass gene list. Approximately 15% were found in at least one abstract (this amounts to 146 lncRNAs), and the most commonly found were paupar, hotair, hottip, neat1, and hoxa-as2. In addition, just over half (53%) of the lncRNAs we found in cancer associated publications were also linked to at least one of 12 tissue types (all tissue types mentioned here except KIRP). This suggests that the non-coding genes the classifier identified have strong tissue specific and cancer associations.

## 3 Discussion

Cancer is a collection of more than two hundred different diseases. The inherent heterogeneity and the presence of many subtypes make treatment difficult. Identifying tumours early, prior to metastatic spread, enables treatment options that lead to better prognosis. Screening approaches for certain cancers (cervical, breast, and colorectal) and early identification of possible neoplastic lesions have helped reduce morbidity and mortality due to those cancers. A fundamental requirement is to distinguish cancer from a non-cancer tissue sample accurately, with high precision and recall. We have utilised epigenetic changes in the DNA methylome together with machine and deep learning approach to develop such a method.

Here, we presented binary and multiclass machine learning models to classify 13 cancer types and corresponding normal tissues. Our approach had a good test set performance for all XGBoost-based models, namely an average accuracy of 0.987 for the binary models and an average accuracy of 0.982 for the multiclass model. The most challenging class in the multiclass model was the normal class, which is unsurprising given that this class is the most heterogeneous, containing normal samples from 13 different tissue types. Depending on the availability of training data this method can be extended to detect hundreds of cancer types.

A number of recent methods have also utilised methylation-based approaches to classify cancer. Hao et al. achieved an accuracy of 0.971 on multiclass classification of four cancer types, using TCGA data [18]. Additionally, an accuracy of 0.961 is obtained by Tang et al., in multiclass classification of 14 tissue types [19]. Liu et al. reaches a ROC AUC of 0.989 with multiclass classification of eight cancer types, where some samples are from TCGA [20]. These recent studies demonstrate that machine learning approaches can be used to classify cancer based on DNA methylation data, even when using a highly-filtered small probe list [21, 20, 19, 18]. Multiple probe selection statistical methods were commonly used, such as LASSO, the moderated t-statistic, Maximum–Relevance-Maximum-Distance, and PCA. For example, Hao et al. uses the moderated t-statistic for pre-screening and then LASSO under a multinomial distribution. The majority of these approaches produce tens of probes, often less than 20, for building a final classification model. These probe lists could potentially be biased by the feature selection methods used. In our approach, we did not significantly restrict the probe list through multiple feature selection models. Instead, we let the XGBoost classification model decide which probes were useful for classification from an input set of around 277,000 features. For the multiclass case, this resulted in a large set PCCs, of size 3,388, that provided us with an interpretable model and an explainable list of genomic loci for further analysis.

We were then able to show that these PCCs can robustly classify cancer when fed into a multiclass deep neural network. The performance on most independent (non-TCGA) data sets was above an *F*_1_ score of 0.8, and half of the independent data sets achieved an *F*_1_ score of over 0.9. These independent data sets were more heterogeneous and reflected more realistic situations. There were two exceptions to the performance of our models, one of them being the independent colon adenocarcinoma (COAD) data set. As indicated in the Results, this low performance can be explained by all adenomas, labelled as normal, being predicted as cancer. The vast majority of colorectal polyps are either adenomas or hyperplastic polyps [22]. Adenomas are dysplastic polyps which can progress via the adenoma-carcinoma sequence to invasive cancer. Therefore, it is common to remove colon adenomas when they are found to stop the possible progression into carcinomas [23, 24]. Thus, this behaviour was inadvertently useful, and when the adenomas were moved from the normal class over to the COAD cancer class, the accuracy increased to 0.897, the MCC to 0.790, and the ROC AUC to 0.980. However, it is important to note that a sample size of nine adenomas is not enough to validate that the model consistently classifies adenomas as cancer.

The other exception is the head and neck squamous cell carcinoma (HNSC) independent data set, which has the lowest performance. HNSC is very heterogeneous, in that it can arise from multiple different tissue sites such as the larynx, tongue, oropharynx and many more. However, the independent HNSC data set only stems from one tissue of origin, the oropharynx. The tissue of origins from the HNSC TCGA data were extremely varied, and only nine samples out of the TCGA data were from the oropharynx (and only six to seven in the training set). In addition, half of the independent HNSC data set was HPV positive (HPV+). HPV+ oropharyngeal squamous cell carcinomas are very different to HPV-oropharyngeal carcinomas [25], and they have distinct methylation patterns [26]. The HPV status of the TCGA data is unknown, but as HPV+ tumours are seen more frequently in the oropharynx than other tissues of origin and only a small fraction of the TCGA data is from the oropharynx, it is likely that a very small fraction of the training data was HPV+. Thus, we were testing on HNSC cancer types with very little training data, which could explain the poor performance. In addition, the independent HNSC data were often misclassified as lung adenocarcinoma (LUAD). A UMAP visualisation of all the TCGA data and the HNSC independent data, as shown in Figure 7, illustrates that out of all of the TCGA classes, the independent HNSC data was the closest to LUAD. This could be due to a biological reason, such as the independent HNSC data are in fact metastases which originated in the lung, or this could be due to a specific data generation or processing artefact that is unique to TCGA LUAD and independent HNSC.

**Figure 7:**
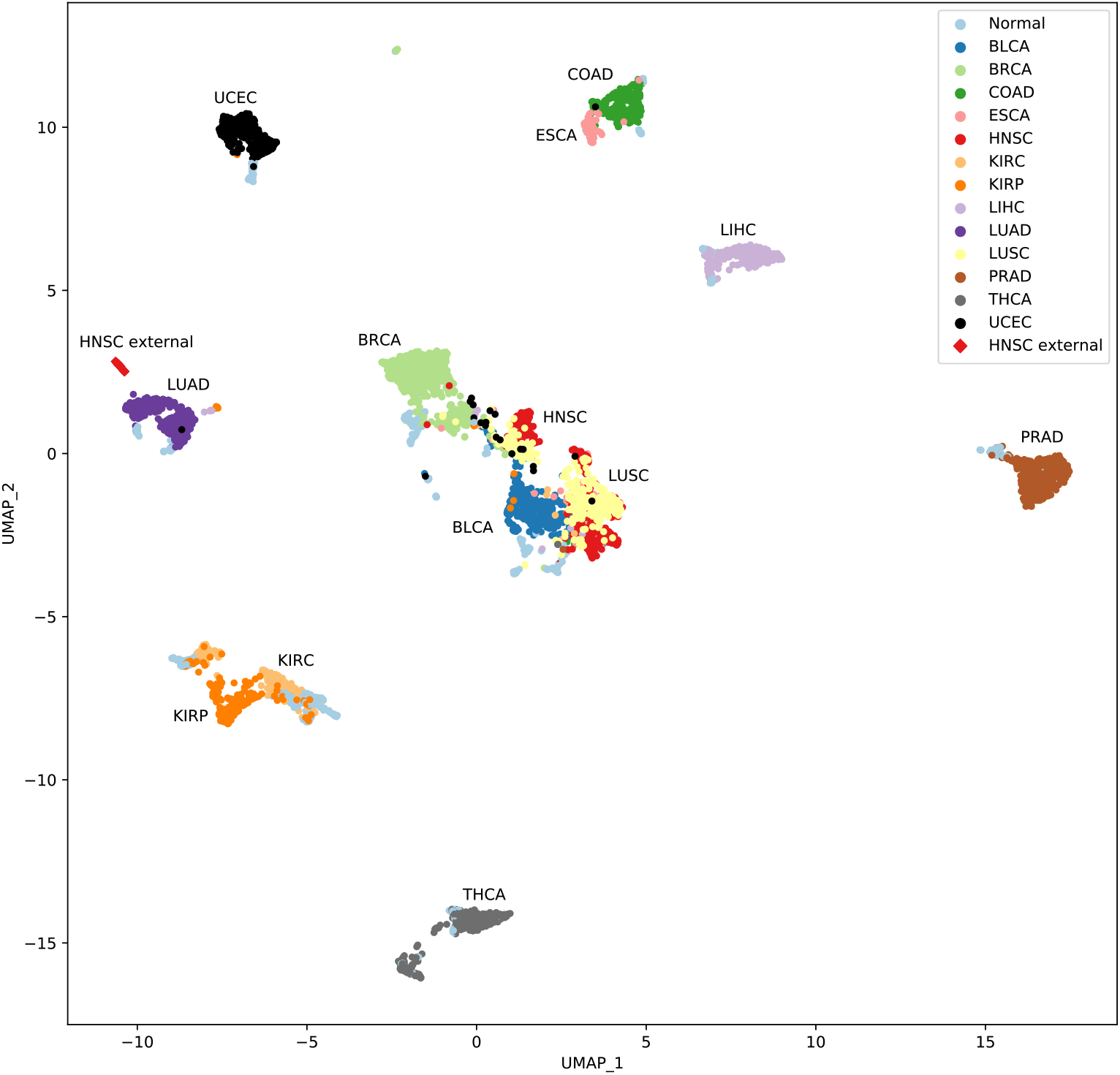
UMAP of all of the TCGA data and the HNSC (head and neck squamous cell carcinoma) independent data set.

Lastly, we showed that the multiclass PCCs do have biologically meaningful significance in cancer. Over-representation analysis revealed that our gene list was enriched in genes important for developmental and differentiation processes, cell communication, metabolic processes, and regulation of transcription. All of these biological processes are linked to cancer hallmarks, and similar methylation studies focusing on various types of cancer also present gene lists enriched in developmental processes [27], metabolic processes [28] and regulation of transcription [29]. Additionally, we showed that our gene lists contain many non-coding RNA genes, primarily consisting of lncRNAs, suggesting that methylation alterations at non-coding gene promoters are frequently present in cancer. This fits with the growing body of research showing that lncRNAs and other non-coding RNAs play a key role in cancer [30, 31, 31].

To conclude, we showed that XGBoost models are suitable for classifying a multitude of cancer types using only DNA methylation data as input. While our multiclass XGBoost model did not perform exceptionally well on independent data sets, we designed a robust neural network that was able to generalise to most independent data sets. In addition, we find that mapping the PCCs to genes produces gene lists that are enriched in functional properties linked to carcinogenesis. Future applications include extending this approach to DNA methylation data of cell-free DNA, with the eventual aim being early detection of multiple types of cancer from liquid biopsy approaches(from blood, cerebrospinal fluid, urine etc.). Furthermore, a clear clinical application of this method is screening for specific cancer types. Although the current models are not optimised for this purpose, achieving the very low false negative rates necessary for screening can be achieved by careful choice of the classification threshold or optimising for recall during model training.

## 4 Methods

### 4.1 Microarray based methylation analysis

Methylome microarray data were obtained from the TCGA GDC data portal^1^. The data sets utilised were from the Illumina Infinium Human DNA Methylation 450 platform, and 13 cancer types with at least 15 normal samples were analysed. Table 3 shows the number of cancer and normal samples for each cancer type.

**Table 3:** Number of normal and cancer samples for each cancer type from TCGA

|  | BLCA | BRCA | COAD | ESCA | HNSC | KIRC | KIRP | LIHC | LUAD | LUSC | PRAD | THCA | UCEC | Total |
| --- | --- | --- | --- | --- | --- | --- | --- | --- | --- | --- | --- | --- | --- | --- |
| Normal | 21 | 96 | 38 | 16 | 50 | 160 | 45 | 50 | 32 | 42 | 50 | 56 | 46 | **702** |
| Cancer | 415 | 793 | 315 | 186 | 530 | 325 | 276 | 380 | 475 | 370 | 503 | 515 | 439 | **5522** |
| Total | 436 | 889 | 353 | 202 | 580 | 485 | 321 | 430 | 507 | 412 | 553 | 571 | 485 | **6224** |

In addition to the TCGA data, a number of data sets from independent studies were also used in model evaluation. All eight independent data sets used were from the Illumina Infinium Human DNA Methylation 450 platform. Where available, raw methylated and unmethylated counts were used to calculate beta values (without any normalisation), but in the case of THCA only normalised beta values were available. The number of cancer and normal samples for each independent data set are shown in Table 4. The sources of each data set are: BRCA: GSE52865, COAD: GSE77955 (only samples from sites colon, left colon, right colon, and sigmoid are taken), ESCA: GSE72874, HNSC: GSE38266 (note that half of these samples are HPV+), KIRC: GSE61441, LIHC: GSE75041, PRAD: project PRAD-CA from ICGC, THCA: GSE97466.

**Table 4:** Number of normal and cancer samples for the independent data sets

|  | BRCA | COAD | ESCA | HNSC | KIRC | LIHC | PRAD | THCA | Total |
| --- | --- | --- | --- | --- | --- | --- | --- | --- | --- |
| Normal | 17 | 20 | 75 | 0 | 46 | 0 | 0 | 67 | **152** |
| Cancer | 40 | 9 | 125 | 42 | 46 | 66 | 199 | 74 | **488** |
| Total | 57 | 29 | 200 | 42 | 92 | 66 | 199 | 141 | **640** |

### 4.2 Data pre-processing

TCGA data was downloaded using the TCGAbiolinks package [33] in R (version 3.6.2) [34], with the following pre-processing steps applied separately for each cancer type. Probes listed as potentially noisy by Naeem et al. [35] were discarded, and only probes mapping to autosomal and sex chromosomes were kept. In addition, we discarded probes with > 5% of missing values. The remaining missing values were imputed using a k-nearest neighbours approach with the impute package in R (k=10, rowmax=0.25) [36]. This resulted in around 277,000 features (probes) per sample (this number varied between cancer types as processing was applied separately to each one). As M-values are more homoscedastic than beta-values [37], we transformed beta-values to M-values using the function:

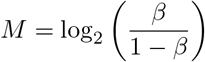

For the multiclass data, features were obtained by taking the intersection of features of every cancer type, and the normal class was obtained by pooling all the normal samples from different tissue types together.

When pre-processing non-TCGA data sets, where the non-normalised data were available, we computed the beta values from methylated and unmethylated counts, and selected the same features as the previously processed TCGA data. Many data sets contained NA values, and unlike before, where we relied on the whole data set to impute the values, we set them to a constant beta of 0.5. Lastly, these beta values were transformed to M-values.

### 4.3 Classification models and metrics

Throughout this study we used both binary and multiclass models. Each binary model compares one tissue type, distinguishing cancer from normal, and the multiclass models utilise all thirteen tissue types and normal samples. Note that in the binary models, the normal class is only normal samples for that tissue, whereas in the multiclass models, the normal class is normal samples from all tissue types pooled together. For each model, the input data were split into training and test sets, with 25% of samples in the test sets.

To begin with, we tested two simple classification models: logistic regression and an SVM. Both models were created and tuned using the package sklearn [38] in Python (version 3.7.5). Hyperparameter tuning on the training set using 5-fold cross-validation selected the default values in most cases, except for the binary logistic regression using the Newton solver and the multiclass SVM using gamma = ‘auto’.

We also created an XGBoost model based on gradient boosted decision trees, using the XGBoost package in Python [16]. Decision trees are constructed in succession so that new trees correct the mistakes of previous trees. Hyperparameter tuning for the binary models resulted in 450 estimators with a maximum depth of 10 and a learning rate of 0.189. The multiclass model has 800 estimators with a max depth of 3 and the same learning rate. In this model, 50% of features are randomly sampled when constructing each tree and 50% of samples are taken in each iteration, which helped to prevent over-fitting.

The final model we used was a multiclass feed forward neural network, which is based on the multiclass XGBoost model. XGBoost models assign an importance to each of their input features, and an importance above zero indicates that this feature is helpful for classification. All input features with an importance greater than zero (3,388 features) were used as input for the neural network. Similar to all other models, we trained it on our TCGA training set. We conducted a hyperparameter search using the Python package Talos [39], using 30% of the training data as a validation set for each hyperparameter test. For training, we used the Adam learning algorithm [40] with cross-entropy loss. A variant of early stopping was used, where the model was trained for a full 500 epochs, and the model at the epoch with the highest validation set accuracy was taken as the final model.

For evaluation, we used standard accuracy, precision and recall metrics. We also report the F_1_ score, which is the harmonic mean of precision and recall:

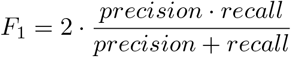

In addition, we report the Matthews Correlation Coefficient (MCC) measure, as it portrays a more comprehensive measure of performance, especially with imbalanced classes in the binary case [41]:

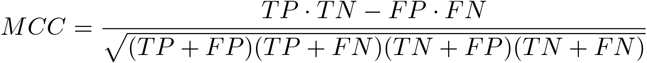

We also report the Area Under the Curve (AUC) for both Receiver Operating Characteristic (ROC) curves and Precision-Recall curves. For both metrics, 1 is a perfect score and 0.5 is the score from a random model (assuming classes are balanced).

### 4.4 Feature Interpretation

#### 4.4.1 Probe annotation and mapping

Probes with an XGBoost importance score > 0 were mapped to the nearest gene, if it overlapped a gene, or was within 5000 base pairs upstream or downstream from a gene’s transcription start or stop site. We mapped each probe to all genes that fulfilled this property. The gene annotation data were obtained from the Ensembl (version 98) using the R package biomaRt [42, 43], and the mapping functionality was implemented using the R package ChIPpeakAnno [44].

#### 4.4.2 Differential methylation analysis

Differential methylation analysis was performed using the R package TCGAbiolinks, and the input data were M-values of the probes after filtering (see Data pre-processing). Differentially methylated probes were found by the Wilcoxon test using the Benjamini-Hochberg adjustment method. Here, a probe had to have an absolute mean difference of above 3 and an adjusted p-value of below 1e-30 to be labelled as differentially methylated.

#### 4.4.3 Gene Ontology Over-Representation Analysis

Functional enrichment analysis was carried out using the R package gprofiler2 [45] with the Bonferroni correction method. The background set were the XGBoost input probes (i.e. the microarray probe list after filtering) mapped to genes. This result was then visualised by REVIGO [46] using the settings: medium, Homo sapiens GO terms, SimRel similarity. The scatter plot in Figure 6**b** was based on the provided R script, and the visible labels were manually selected to provide information across each cluster and focus on terms with low adjusted p-values.

## 5 Data Availability

The results shown here are in whole or part based upon methylome data generated by the TCGA Research Network: https://www.cancer.gov/tcga. Non TCGA evaluation data sets with the following accession IDs were downloaded from NCBI GEO and ICGC.

|  |  |
| --- | --- |
| BRCA | GSE52865 |
| COAD | GSE77955 |
| ESCA | GSE72874 |
| HNSC | GSE38266 |
| KIRC | GSE61441 |
| LIHC | GSE75041 |
| PRAD | PRAD-CA from ICGC |
| THCA | GSE97466 |

## 6 Reproducible research

The code for this project was produced in a reproducible manner with Python Jupyter and R notebooks, and will be made publicly available on GitHub after peer-reviewed publication.

## 7 Author Contribution

S.A.S and C.M conceived the study. I.N developed the machine learning models and carried out the data processing and analysis. M.S contributed to the initial analysis during a summer studentship. C.M, S.J and R.F contributed data sets and provided guidance. I.N and S.A.S wrote the manuscript with input from the other authors.

## 8 Acknowledgements

S.A.S, I.N, and M.S were supported by the UK Medical Research Council core funding (MC_UU_12022/10) to the S.A.S laboratory. We also thank members of the S.A.S laboratory that read and commented on the manuscript.

## 9 Abbreviations

TCGA: The Cancer Genome Atlas
BLCA: Bladder urothelial carcinoma
BRCA: Breast invasive carcinoma
COAD: Colon adenocarcinoma
ESCA: Esophageal carcinoma
HNSC: Head and neck squamous cell carcinoma
KIRC: Kidney renal clear cell carcinoma
KIRP: Kidney renal papillary cell carcinoma
LIHC: Liver hepatocellular carcinoma
LUAD: Lung adenocarcinoma
LUSC: Lung squamous cell carcinoma
PRAD: Prostate adenocarcinoma
THCA: Thyroid carcinoma
UCEC: Uterine corpus endometrial carcinoma
AUC: Area Under the Curve
ROC: Receiver Operating Characteristic
MCC: Matthews Correlation Coefficient
UMAP: Uniform manifold approximation and projection

## Footnotes

1 https://portal.gdc.cancer.gov/

