## Supplementary Tables S1-5 for "Machine Classification of Methylomes in Cancer"

**S3 Metric tables for XGBoost binary models** **5**

**S4 Metric table for neural network model on TCGA testset** **6**

**S5 Metrics table for multiclass XGBoost on external data** **6**

**Abbreviations**

| | |
|---|---|
| BLCA | Bladder Urothelial Carcinoma |
| BRCA | Breast invasive carcinoma |
| COAD | Colon adenocarcinoma |
| ESCA | Esophageal carcinoma |
| HNSC | Head and Neck squamous cell carcinoma |
| KIRC | Kidney renal clear cell carcinoma |
| KIRP | Kidney renal papillary cell carcinoma |
| LIHC | Liver hepatocellular carcinoma |
| LUAD | Lung adenocarcinoma |
| LUSC | Lung squamous cell carcinoma |
| PRAD | Prostate adenocarcinoma |
| THCA | Thyroid carcinoma |
| UCEC | Uterine Corpus Endometrial Carcinoma |

### S1 Metric tables for Logistic Regression models

#### S1.1 Binary

**a. BLCA**

| | Normal | BLCA |
|---|---|---|
| Precision | 1.0 | 1.0 |
| Recall | 1.0 | 1.0 |
| $F_1$ | 1.0 | 1.0 |
| Accuracy | 1.0 | |
| MCC | 1.0 | |

**b. BRCA**

| | Normal | BRCA |
|---|---|---|
| Precision | 0.957 | 0.99 |
| Recall | 0.917 | 0.995 |
| $F_1$ | 0.936 | 0.992 |
| Accuracy | 0.987 | |
| MCC | 0.929 | |

**c. COAD**

| | Normal | COAD |
|---|---|---|
| Precision | 1.0 | 1.0 |
| Recall | 1.0 | 1.0 |
| $F_1$ | 1.0 | 1.0 |
| Accuracy | 1.0 | |
| MCC | 1.0 | |

**d. ESCA**

| | Normal | ESCA |
|---|---|---|
| Precision | 1.0 | 1.0 |
| Recall | 1.0 | 1.0 |
| $F_1$ | 1.0 | 1.0 |
| Accuracy | 1.0 | |
| MCC | 1.0 | |

**e. HNSC**

| | Normal | HNSC |
|---|---|---|
| Precision | 1.0 | 0.992 |
| Recall | 0.923 | 1.0 |
| $F_1$ | 0.96 | 0.996 |
| Accuracy | 0.993 | |
| MCC | 0.957 | |

**f. KIRC**

| | Normal | KIRC |
|---|---|---|
| Precision | 1.0 | 1.0 |
| Recall | 1.0 | 1.0 |
| $F_1$ | 1.0 | 1.0 |
| Accuracy | 1.0 | |
| MCC | 1.0 | |

**g. KIRP**

| | Normal | KIRP |
|---|---|---|
| Precision | 0.917 | 1.0 |
| Recall | 1.0 | 0.986 |
| $F_1$ | 0.957 | 0.993 |
| Accuracy | 0.988 | |
| MCC | 0.951 | |

**h. LIHC**

| | Normal | LIHC |
|---|---|---|
| Precision | 0.929 | 1.0 |
| Recall | 1.0 | 0.989 |
| $F_1$ | 0.963 | 0.995 |
| Accuracy | 0.991 | |
| MCC | 0.959 | |

**i. LUAD**

| | Normal | LUAD |
|---|---|---|
| Precision | 0.889 | 1.0 |
| Recall | 1.0 | 0.992 |
| $F_1$ | 0.941 | 0.996 |
| Accuracy | 0.992 | |
| MCC | 0.939 | |

**j. LUSC**

| | Normal | LUSC |
|---|---|---|
| Precision | 0.917 | 1.0 |
| Recall | 1.0 | 0.989 |
| $F_1$ | 0.957 | 0.995 |
| Accuracy | 0.99 | |
| MCC | 0.952 | |

**k. PRAD**

| | Normal | PRAD |
|---|---|---|
| Precision | 1.0 | 0.992 |
| Recall | 0.923 | 1.0 |
| $F_1$ | 0.96 | 0.996 |
| Accuracy | 0.993 | |
| MCC | 0.957 | |

**l. THCA**

| | Normal | THCA |
|---|---|---|
| Precision | 0.737 | 1.0 |
| Recall | 1.0 | 0.961 |
| $F_1$ | 0.848 | 0.98 |
| Accuracy | 0.965 | |
| MCC | 0.842 | |

**m. UCEC**

| | Normal | UCEC |
|---|---|---|
| Precision | 1.0 | 1.0 |
| Recall | 1.0 | 1.0 |
| $F_1$ | 1.0 | 1.0 |
| Accuracy | 1.0 | |
| MCC | 1.0 | |

##### S1.2 Multiclass

| | Normal | BLCA | BRCA | COAD | ESCA | HNSC | KIRC | KIRP | LIHC | LUAD | LUSC | PRAD | THCA | UCEC |
|---|---|---|---|---|---|---|---|---|---|---|---|---|---|---|
| precision | 0.953 | 0.99 | 0.98 | 0.988 | 0.956 | 0.949 | 0.975 | 0.957 | 1.0 | 0.983 | 0.977 | 0.954 | 0.992 | 1.0 |
| recall | 0.937 | 0.99 | 0.99 | 1.0 | 0.935 | 0.977 | 0.963 | 0.957 | 0.968 | 1.0 | 0.914 | 0.984 | 1.0 | 1.0 |
| f1 | 0.945 | 0.99 | 0.985 | 0.994 | 0.945 | 0.963 | 0.969 | 0.957 | 0.984 | 0.992 | 0.944 | 0.969 | 0.996 | 1.0 |
| accuracy | | | | | | | 0.975 | | | | | | | |
| mcc | | | | | | | 0.973 | | | | | | | |

### S2 Metrics tables for Support Vector Machine models

#### S2.1 Binary

**a. BLCA**

| | Normal | BLCA |
|---|---|---|
| Precision | 1.0 | 1.0 |
| Recall | 1.0 | 1.0 |
| $F_1$ | 1.0 | 1.0 |
| Accuracy | 1.0 | |
| MCC | 1.0 | |

**b. BRCA**

| | Normal | BRCA |
|---|---|---|
| Precision | 0.957 | 0.99 |
| Recall | 0.917 | 0.995 |
| $F_1$ | 0.936 | 0.992 |
| Accuracy | 0.987 | |
| MCC | 0.929 | |

**c. COAD**

| | Normal | COAD |
|---|---|---|
| Precision | 1.0 | 1.0 |
| Recall | 1.0 | 1.0 |
| $F_1$ | 1.0 | 1.0 |
| Accuracy | 1.0 | |
| MCC | 1.0 | |

**d. ESCA**

| | Normal | ESCA |
|---|---|---|
| Precision | 1.0 | 1.0 |
| Recall | 1.0 | 1.0 |
| $F_1$ | 1.0 | 1.0 |
| Accuracy | 1.0 | |
| MCC | 1.0 | |

**e. HNSC**

| | Normal | HNSC |
|---|---|---|
| Precision | 1.0 | 0.992 |
| Recall | 0.923 | 1.0 |
| $F_1$ | 0.96 | 0.996 |
| Accuracy | 0.993 | |
| MCC | 0.957 | |

**f. KIRC**

| | Normal | KIRC |
|---|---|---|
| Precision | 1.0 | 1.0 |
| Recall | 1.0 | 1.0 |
| $F_1$ | 1.0 | 1.0 |
| Accuracy | 1.0 | |
| MCC | 1.0 | |

**g. KIRP**

| | Normal | KIRP |
|---|---|---|
| Precision | 1.0 | 1.0 |
| Recall | 1.0 | 1.0 |
| $F_1$ | 1.0 | 1.0 |
| Accuracy | 1.0 | |
| MCC | 1.0 | |

**h. LIHC**

| | Normal | LIHC |
|---|---|---|
| Precision | 0.929 | 1.0 |
| Recall | 1.0 | 0.989 |
| $F_1$ | 0.963 | 0.995 |
| Accuracy | 0.991 | |
| MCC | 0.959 | |

**i. LUAD**

| | Normal | LUAD |
|---|---|---|
| Precision | 0.889 | 1.0 |
| Recall | 1.0 | 0.992 |
| $F_1$ | 0.941 | 0.996 |
| Accuracy | 0.992 | |
| MCC | 0.939 | |

**j. LUSC**

| | Normal | LUSC |
|---|---|---|
| Precision | 0.917 | 1.0 |
| Recall | 1.0 | 0.989 |
| $F_1$ | 0.957 | 0.995 |
| Accuracy | 0.99 | |
| MCC | 0.952 | |

**k. PRAD**

| | Normal | PRAD |
|---|---|---|
| Precision | 1.0 | 0.992 |
| Recall | 0.923 | 1.0 |
| $F_1$ | 0.96 | 0.996 |
| Accuracy | 0.993 | |
| MCC | 0.957 | |

**l. THCA**

| | Normal | THCA |
|---|---|---|
| Precision | 0.778 | 1.0 |
| Recall | 1.0 | 0.969 |
| $F_1$ | 0.875 | 0.984 |
| Accuracy | 0.972 | |
| MCC | 0.868 | |

**m. UCEC**

| | Normal | UCEC |
|---|---|---|
| Precision | 1.0 | 1.0 |
| Recall | 1.0 | 1.0 |
| $F_1$ | 1.0 | 1.0 |
| Accuracy | 1.0 | |
| MCC | 1.0 | |

##### S2.2 Multiclass

| | Normal | BLCA | BRCA | COAD | ESCA | HNSC | KIRC | KIRP | LIHC | LUAD | LUSC | PRAD | THCA | UCEC |
|---|---|---|---|---|---|---|---|---|---|---|---|---|---|---|
| Precision | 0.915 | 0.953 | 0.975 | 1.0 | 0.941 | 0.899 | 0.975 | 0.971 | 1.0 | 1.0 | 0.925 | 0.946 | 0.977 | 0.991 |
| Recall | 0.926 | 0.981 | 0.99 | 1.0 | 0.696 | 0.947 | 0.963 | 0.957 | 0.937 | 0.992 | 0.925 | 0.968 | 0.992 | 1.0 |
| $F_1$ | 0.92 | 0.967 | 0.982 | 1.0 | 0.8 | 0.923 | 0.969 | 0.964 | 0.967 | 0.996 | 0.925 | 0.957 | 0.985 | 0.995 |
| Accuracy | 0.96 | | | | | | | | | | | | | |
| MCC | 0.956 | | | | | | | | | | | | | |

#### S3 Metric tables for XGBoost binary models

###### a. BLCA

| | Normal | BLCA |
|---|---|---|
| Precision | 1.0 | 0.981 |
| Recall | 0.6 | 1.0 |
| $F_1$ | 0.75 | 0.99 |
| Accuracy | 0.982 | |
| MCC | 0.767 | |

###### b. BRCA

| | Normal | BRCA |
|---|---|---|
| Precision | 1.0 | 0.99 |
| Recall | 0.917 | 1.0 |
| $F_1$ | 0.957 | 0.995 |
| Accuracy | 0.991 | |
| MCC | 0.953 | |

###### c. COAD

| | Normal | COAD |
|---|---|---|
| Precision | 1.0 | 1.0 |
| Recall | 1.0 | 1.0 |
| $F_1$ | 1.0 | 1.0 |
| Accuracy | 1.0 | |
| MCC | 1.0 | |

###### d. ESCA

| | Normal | ESCA |
|---|---|---|
| Precision | 1.0 | 0.959 |
| Recall | 0.5 | 1.0 |
| $F_1$ | 0.667 | 0.979 |
| Accuracy | 0.961 | |
| MCC | 0.693 | |

###### e. HNSC

| | Normal | HNSC |
|---|---|---|
| Precision | 0.923 | 1.0 |
| Recall | 1.0 | 0.992 |
| $F_1$ | 0.96 | 0.996 |
| Accuracy | 0.993 | |
| MCC | 0.957 | |

###### f. KIRC

| | Normal | KIRC |
|---|---|---|
| Precision | 1.0 | 1.0 |
| Recall | 1.0 | 1.0 |
| $F_1$ | 1.0 | 1.0 |
| Accuracy | 1.0 | |
| MCC | 1.0 | |

###### g. KIRP

| | Normal | KIRP |
|---|---|---|
| Precision | 0.833 | 0.986 |
| Recall | 0.909 | 0.971 |
| $F_1$ | 0.87 | 0.978 |
| Accuracy | 0.963 | |
| MCC | 0.849 | |

###### h. LIHC

| | Normal | LIHC |
|---|---|---|
| Precision | 1.0 | 0.99 |
| Recall | 0.923 | 1.0 |
| $F_1$ | 0.96 | 0.995 |
| Accuracy | 0.991 | |
| MCC | 0.956 | |

**i. LUAD**

| | Normal | LUAD |
|---|---|---|
| Precision | 1.0 | 1.0 |
| Recall | 1.0 | 1.0 |
| $F_1$ | 1.0 | 1.0 |
| Accuracy | 1.0 | |
| MCC | 1.0 | |

**j. LUSC**

| | Normal | LUSC |
|---|---|---|
| Precision | 1.0 | 1.0 |
| Recall | 1.0 | 1.0 |
| $F_1$ | 1.0 | 1.0 |
| Accuracy | 1.0 | |
| MCC | 1.0 | |

**k. PRAD**

| | Normal | PRAD |
|---|---|---|
| Precision | 0.75 | 0.992 |
| Recall | 0.923 | 0.968 |
| $F_1$ | 0.828 | 0.98 |
| Accuracy | 0.964 | |
| MCC | 0.813 | |

**l. THCA**

| | Normal | THCA |
|---|---|---|
| Precision | 1.0 | 0.992 |
| Recall | 0.929 | 1.0 |
| $F_1$ | 0.963 | 0.996 |
| Accuracy | 0.993 | |
| MCC | 0.96 | |

**m. UCEC**

| | Normal | UCEC |
|---|---|---|
| Precision | 1.0 | 1.0 |
| Recall | 1.0 | 1.0 |
| $F_1$ | 1.0 | 1.0 |
| Accuracy | 1.0 | |
| MCC | 1.0 | |

### S4 Metric table for neural network model on TCGA testset

| | Normal | BLCA | BRCA | COAD | ESCA | HNSC | KIRC | KIRP | LIHC | LUAD | LUSC | PRAD | THCA | UCEC |
|---|---|---|---|---|---|---|---|---|---|---|---|---|---|---|
| Precision | 0.958 | 0.99 | 0.98 | 0.988 | 0.978 | 0.963 | 0.963 | 0.957 | 1.0 | 1.0 | 1.0 | 0.947 | 0.985 | 1.0 |
| Recall | 0.92 | 0.99 | 0.98 | 1.0 | 0.978 | 0.992 | 0.963 | 0.971 | 0.979 | 1.0 | 0.957 | 0.984 | 1.0 | 1.0 |
| $F_1$ | 0.939 | 0.99 | 0.98 | 0.994 | 0.978 | 0.978 | 0.963 | 0.964 | 0.989 | 1.0 | 0.978 | 0.965 | 0.992 | 1.0 |
| Accuracy | 0.978 | | | | | | | | | | | | | |
| MCC | 0.976 | | | | | | | | | | | | | |

### S5 Metrics table for multiclass XGBoost on external data

| | Normal | BRCA | COAD | ESCA | HNSC | KIRC | LIHC | PRAD | THCA |
|---|---|---|---|---|---|---|---|---|---|
| Precision | 0.914 | 0.971 | 0.333 | 0.893 | 1.0 | 1.0 | 1.0 | 0.98 | 0.861 |
| Recall | 0.8 | 0.85 | 0.778 | 0.936 | 0.167 | 0.348 | 1.0 | 0.251 | 0.838 |
| $F_1$ | 0.853 | 0.907 | 0.467 | 0.914 | 0.286 | 0.516 | 1.0 | 0.4 | 0.849 |
| Accuracy | 0.653 | | | | | | | | |
| MCC | 0.645 | | | | | | | | |
